# Regulatory Role for Tumor Suppressor REST on Estrogen Receptor (*ESR1*) Expression and Leiomyoma Pathophysiology

**DOI:** 10.1101/2025.08.29.673140

**Authors:** Skylar G. Bird, Sumedha Gunewardena, Ashley Cloud, Sornakala Ganeshkumar, Vargheese M. Chennathukuzhi

**Author notes:** Correspondence: Vargheese M. Chennathukuzhi, 3089 HLSIC, 3901 Rainbow Blvd, Kansas City, Kansas 66160, USA. **Author contributions** VMC designed the research; SGB, AC, and SG performed the research; SGB and SG analyzed the data; SGB wrote and VMC edited the manuscript.

## Abstract

Uterine fibroids, benign tumors of the smooth muscle layer of the uterus, plague approximately 80% of the female population by age 50. While there have been efforts to understand the mechanism behind this pathophysiology, it largely remains unclear. Lack of preclinical animal models that recapitulate aberrant steroid hormone pathways in UL has significantly hampered the development of long-term hormonal therapies for uterine fibroids. In addition, cultured myometrial as well as leiomyoma smooth muscle cells rapidly silence both estrogen receptor alpha (*ESR1*) and progesterone receptor (*PGR*) expression through unknown mechanisms, further limiting *in vitro* mechanistic studies of UL. Previous work by our lab has determined the loss of REST, a master regulator of epigenetic gene silencing, in leiomyoma results in the upregulation of ESR1 targets and therefore estrogen signaling. Using ChIP-PCR, we find REST is directly associated with *ESR1* genomic locus, playing a role in its epigenetic regulation. ChIP-seq analysis of *Rest* cKO mouse uterus samples reveals a global role for REST in the regulation of progesterone receptor target genes and highlights alterations in PGR binding within the *Esr1* locus. Additionally, we find REST inhibition of ESR1 expression is regulated through upstream WNT planar cell polarity molecule, PRICKLE1. Based on role of REST in silencing ESR1 expression in cultured myometrial cells, our results support the development of a potential cell culture method to maintain ESR1 expression through REST modulation. Finally, we establish a broad role for REST in epigenetic regulation relevant to leiomyoma pathophysiology.

## Introduction

Uterine fibroids, otherwise known as leiomyomas, can cost the United States upwards of 30 billion dollars annually in medical care [1, 2]. These benign neoplasms grow and develop in the smooth muscle layer (myometrium) of the uterus. Leiomyomas require women, often pre-peak reproductive age, to undergo life-changing medical interventions such as hysterectomy or myomectomy [3]. These surgeries result in both lifelong physical and psychological consequences altering or ending the patient’s ability to conceive [4, 5]. In addition, leiomyomas are known to impact African American women at a higher incidence rate with a more severe prognosis [6-8]. There is yet to be a well-established explanation for this racial disparity. Pharmacological treatments for leiomyomas are lacking, stressing the need for better non-surgical interventions [9]. This is due to an unclear understanding of leiomyoma pathophysiology and long-term safety concerns of current treatments [10-12].

Estrogen plays a major role in leiomyoma formation through the activation of ESR1 signaling and through the induction of PGR [13]. Leiomyomas are deemed estrogen dependent because they fail to develop before puberty and often reduce in size once a woman begins menopause [13]. Elevated PGR expression induced by elevated ESR1 creates a microenvironment which supports tumor proliferation. In addition, *ESR1* promoter in leiomyomas have epigenetic differences which in turn affect estrogen signaling and tumor development [13]. Furthermore, epigenetic differences accompany atypical ESR1 phosphorylation resulting in atypical phosphorylation of MAPK [13, 14]. Leiomyomas have been found to demonstrate histone epigenomic alterations including changes in H3K27Me3 and H3K4Me3 levels [7].

Additionally, the loss of REST plays a critical role in the development of leiomyomas and enhanced estrogen response [15, 16]. REST, known as RE1-silencing transcription factor or NRSF (neuron-restrictive silencing factor), acts as a transcriptional repressor silencing several neuronal genes and suppressing tumorigenesis [17-20]. REST acts as a transcriptional repressor by binding repressor element 1 (*RE-1*) sites within the regulatory regions of its target genes [19, 21]. Collectively, we found the loss of *Rest* in cKO mice led to the development of a leiomyoma phenotype and activation of the estrogen signaling pathway [15].

Despite this progress, the mechanism by which REST regulates ESR1 and activates estrogen signaling in leiomyoma is unknown. In this study, we demonstrate the loss of REST leads to upregulation of ESR1 expression *in vitro* and *in vivo*. We establish a pathway, which informs a novel cell culture method to maintain ESR1 expression using modulation of levels of REST. We further determine WNT planar cell polarity molecule, PRICKLE1, acts upstream to regulate REST’s inhibition of ESR1. In UL tissue and cultured primary cells, we find that REST directly associates with *ESR1* regulatory region, playing a role in its epigenetic control. Finally, we investigate the global role of REST on mouse uterine gene epigenetic regulation and reveal changes in binding enrichment surrounding the *Esr1* promoter. These findings provide new insight necessary for better therapeutic development based on leiomyoma specific mechanisms of steroid hormone signaling.

## Results

### Loss of REST Leads to Increased *ESR1* Expression in Leiomyoma

Previously, we showed the loss of *Rest* in uterine cKO mouse models resulted in significant activation of estrogen receptor signaling [15]. To investigate the effect of REST loss in UL, we quantified RNA levels of *ESR1* derived from matched myometrium and leiomyoma samples from patients undergoing hysterectomy. Quantitative RT-qPCR confirmed a significant increase in *ESR1* expression in leiomyoma samples compared to myometrium (Figure 1A). In addition, we investigated *Esr1* levels in our *Rest*^*f/f*^*PR*^*+/Cre*^ mouse model where *Rest* is lost in all layers of the uterus. RNA sequencing revealed a significant increase in *Esr1* expression in all three layers of the uterus in the cKO compared to the wildtype (Figure 1B). Additionally, immunofluorescence staining confirmed a significant increase in ESR1 expression in our *Rest* cKO mouse model, Rest^f/f^Amhr2^+/Cre^, where *Rest* gene is ablated in mesenchymal cells compared to the wildtype (Figure 1C).

**Figure 1:**
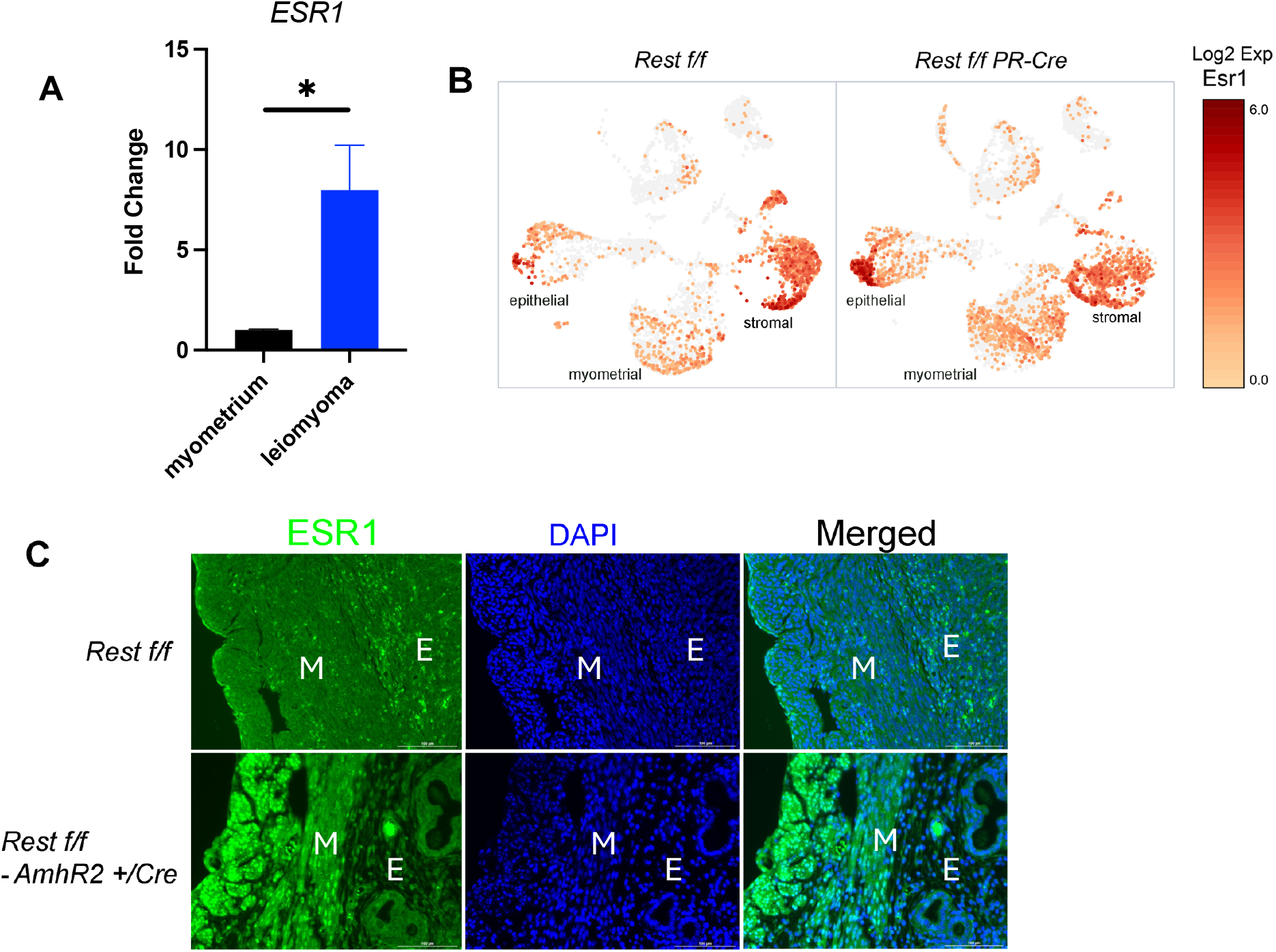
Loss of REST Leads to Increased *ESR1* Expression in Leiomyoma. **(A)** Taqman RT-qPCR of *ESR1* from myometrium and leiomyoma matched patient tissue. **(B)** RNA sequencing investigating *Esr1* expression in wildtype compared to *Rest*^*f/f*^*PR*^*+/Cre*^ cKO mouse model. **(C)** Immunofluorescence staining of ESR1 in wildtype compared to *Rest*^*f/f*^*Amhr2*^*+/Cre*^ cKO mouse uterus at diestrus. M, myometrium, E, endometrial stroma.

### Knockdown of *REST* in Cell Culture Can Restore ESR1 Expression

Myometrial and leiomyoma smooth muscle cells are known to lose estrogen receptor expression once they are cultured. This results in an absolute lack of reliable *in vitro* models to study uterine fibroids. We wanted to investigate if we could create a cell culture model which would restore expression of ESR1. Using an siRNA knockdown system, we knocked down *REST* in primary myometrial cells from passage 1 to 3. REST protein was significantly knocked down for passages one through three (Figure 2A). ESR1 showed a significant increase in protein level for passages two and three (Figure 2A). Furthermore, we analyzed REST and ESR1 protein levels in an immortalized leiomyoma (DDHLM) cell line after lentivirus mediated CRISPR-Cas9 knockout of *REST*. Along with an expected loss of REST protein expression, we observed a significant increase in ESR1 protein levels in the immortalized, CRISPR-edited cell line (Supplemental Figure 1). We performed luciferase assays in cultured myometrial cells to determine the function of ESR1 under *siREST* knockdown conditions. We observed a significant increase in fold change of luciferase activity in myometrial cells with *REST* knocked down when treated with estrogen (Figure 2D). MCF7 cells, used as a positive control since ESR1 response is well established, showed a significant increase in fold change of luciferase activity when treated with estrogen (Figure 2D).

**Figure 2:**
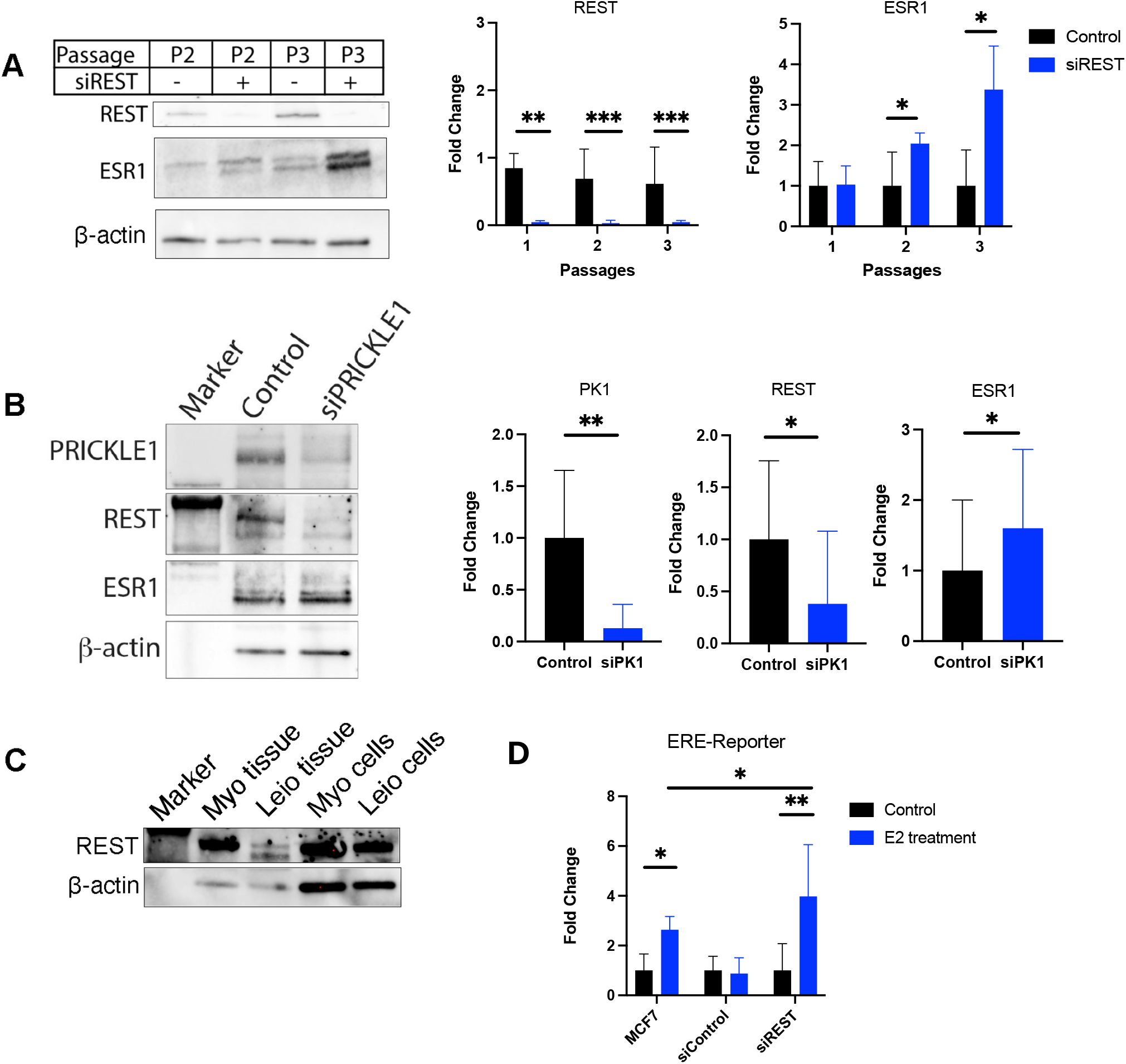
Knockdown of *REST* in Cell Culture Can Restore ESR1 Expression. **(A)** SiRNA knockdown of *REST* in primary myometrial cells western blot and quantification. **(B)** SiRNA knockdown of *PRICKLE1 (PK1*) in primary myometrial cells western blot and quantification. **(C)** Western blot comparing REST levels in myometrium versus leiomyoma tissue and cultured myometrial cells versus cultured leiomyoma cells. (D) Luciferase assay demonstrating functionality of ESR1 after *siREST* knockdown in primary cells.

Previously, we established a role for PRICKLE1 in the upstream regulation of REST. In leiomyoma, PRICKLE1 is lost leading to the loss of REST downstream [22]. We hypothesized PRICKLE1 may be playing a role in the mechanism behind REST’s regulation of ESR1 protein levels. We performed a siRNA knockdown of *PRICKLE1* in primary myometrial cells and quantified PRICKLE1, REST, and ESR1 protein levels by western blot. As expected, *PRICKLE1* knockdown resulted in a significant downregulation of REST protein levels (Figure 2B). In addition, we observed a significant increase in ESR1 protein levels suggesting PRICKLE1 is in fact acting through REST to regulate ESR1 expression (Figure 2B).

### REST Levels Return in Cultured Leiomyoma Cells

We wanted to further investigate if the regulation of REST on ESR1 was being lost in cultured leiomyoma cells compared to leiomyoma tissue, resulting in the silencing of ESR1 *in vitro*. We hypothesized that the loss of REST seen in UL *in vivo* would not be present when the smooth muscle cells are cultured in vitro and restoring low REST levels in cultured leiomyoma cells could maintain ESR1 expression *in vitro*. We analyzed REST protein levels in myometrial tissue and cultured myometrial cells from patients. This was similarly repeated for leiomyoma tissue and cultured cells. Interestingly, lower levels of REST present in leiomyoma tissue were not maintained when the cells were cultured (Figure 2C). This finding, robust expression of REST in leiomyoma cells when cultured, suggests a potential mechanism for loss of estrogen receptor expression *in vitro*.

### Role of REST on Uterine Gene Regulation

Previously, our lab performed ChIP-PCR analysis of several REST-PGR target genes in a *Rest*^*fl/fl*^ *PR*^*+/Cre*^ mouse model [15]. In order to better understand the scope of REST on ESR1 signaling and UL pathogenesis, we investigated the global epigenetic role of REST on uterine gene regulation by ChIP-seq analysis (GSE306878). The distribution of differential binding events of REST in *Rest*^*fl/fl*^ and *Rest*^*fl/fl*^ *PR*^*+/Cre*^ is given in Figure 3A. *Rest*^*fl/fl*^ constituted the highest number of unique REST binding sites (64.25%) followed by *Rest*^*fl/fl*^ *PR*^*+/Cre*^ (25.7%) and sites present in both (10.04%). The distribution of these sites around a 3 kb region of gene TSS was similarly highest for *Rest*^*fl/fl*^ followed by *Rest*^*fl/fl*^ *PR*^*+/Cre*^ (Figure 4B). The distribution of differential binding events of PGR in *Rest*^*fl/fl*^ and *Rest*^*fl/fl*^ *PR*^*+/Cre*^ is given in Figure 3A. *Rest*^*fl/fl*^ constituted the highest number of unique PGR binding sites (75.6%) followed by (23.2%) and sites present in both (1.05%). The distribution heatmap of PGR binding sites around a 3 kb region of gene TSS was enriched in *Rest*^*fl/fl*^ *PR*^*+/Cre*^ showing altered PGR function in the absence of REST. A vast majority of the peaks which passed the cutoff criteria for the PGR antibody contained consensus *Pgre* sequence in *Rest*^*fl/fl*^ and *Rest*^*fl/fl*^ *PR*^*+/Cre*^ (Table 3). Under the REST antibody condition, approximately a quarter of the peaks which passed the cutoff criteria possessed the *RE-1* consensus sequence in *Rest*^*fl/fl*^ and *Rest*^*fl/fl*^ *PR*^*+/Cre*^(Table 3). REST also may be binding globally through the mediation of other transcription factors.

**Figure 3:**
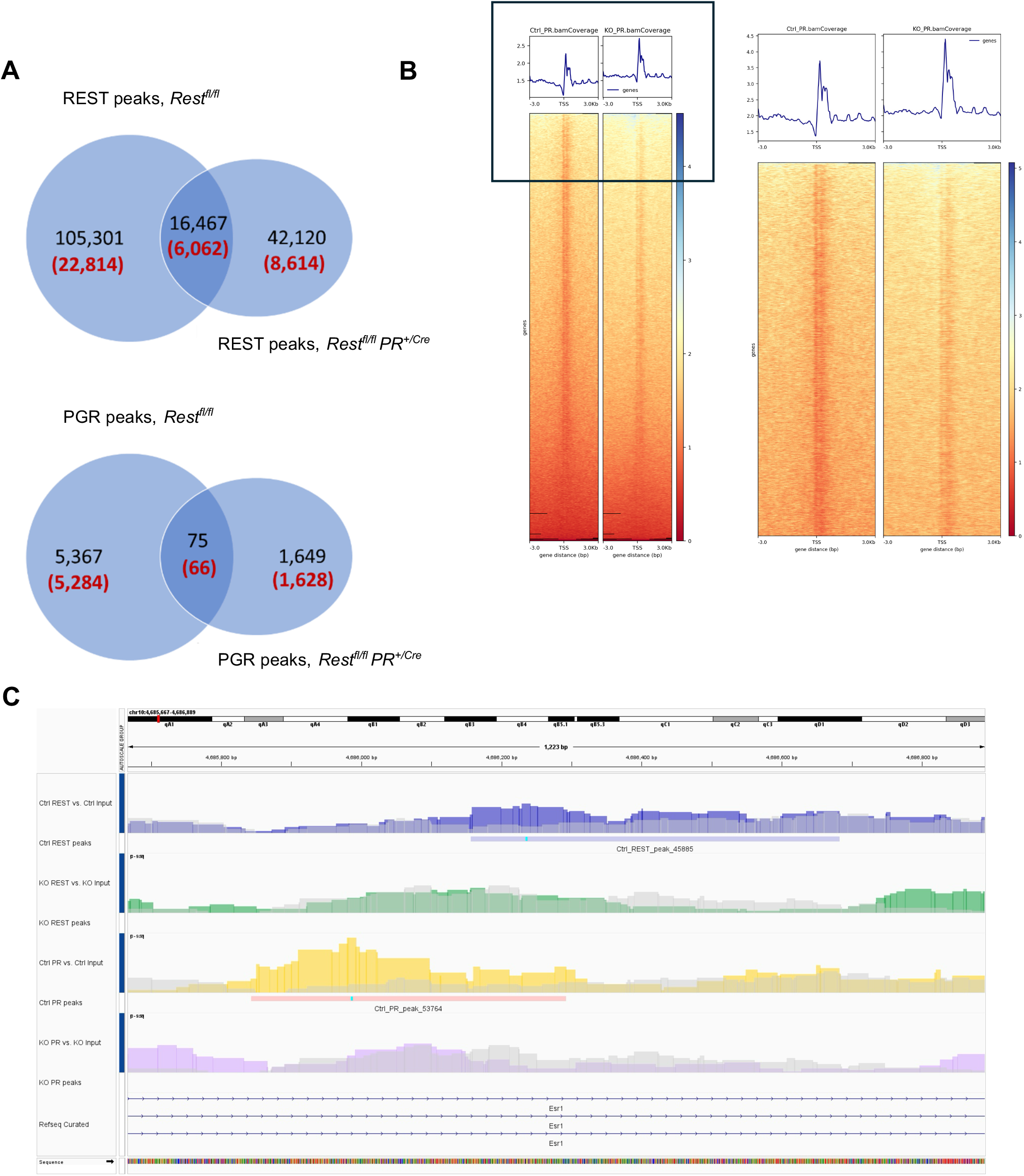
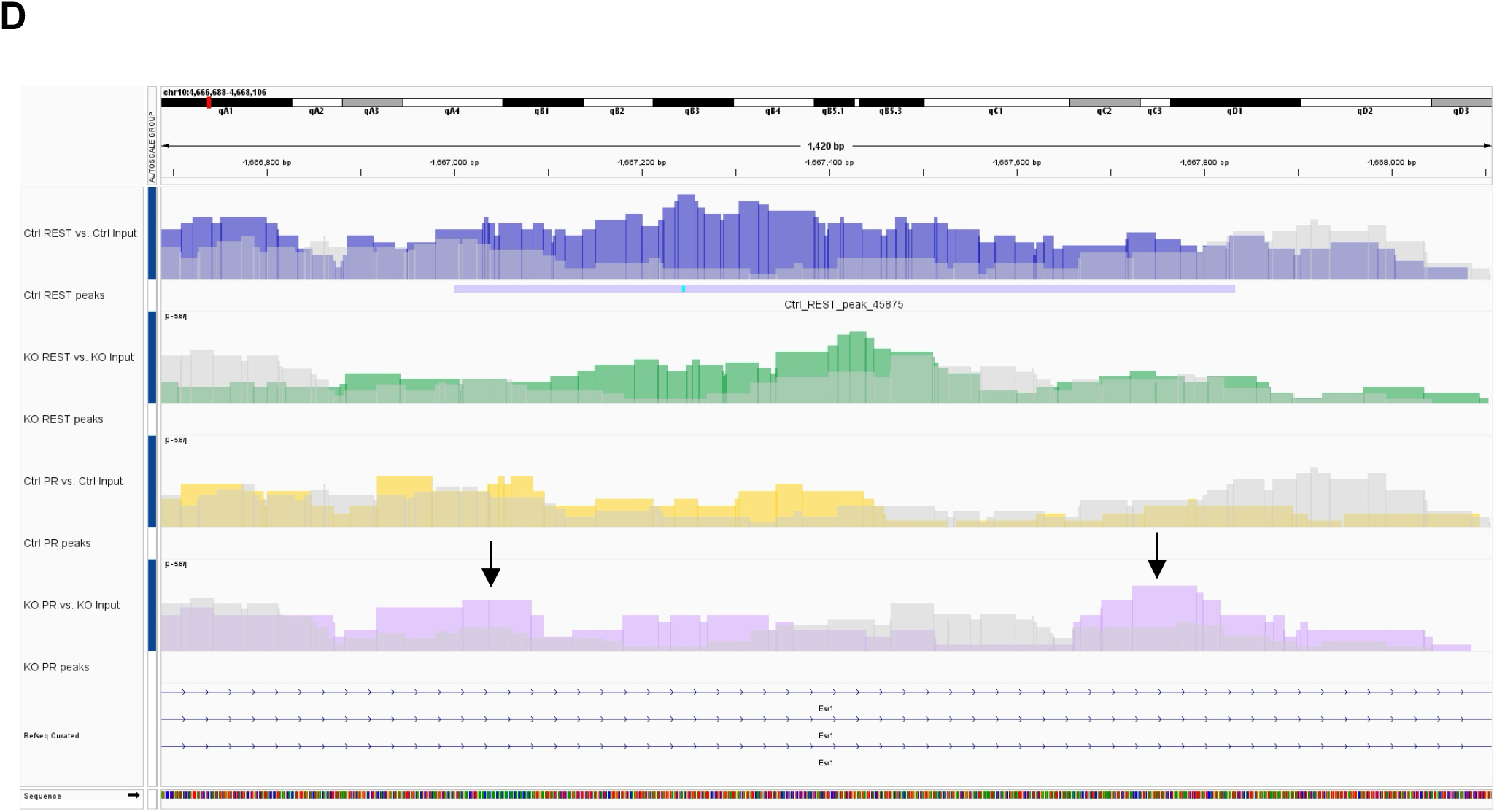
Role of REST on Uterine Gene Regulation. **(A)** Distribution of differential binding events for REST and PGR in *Rest*^*fl/fl*^ and *Rest*^*fl/fl*^ *PR*^*+/Cre*^. **(B)** Heatmap of PGR binding of all genes in control (*Rest*^*fl/fl*^) and KO (*Rest*^*fl/fl*^ *PR*^*+/Cre*^) around TSS. Inset shows enlarged image of top 50 reads. **(C)** Overlay of PGR binding site dependent on REST binding on *Esr1* locus. **(D)** Overlay of PGR binding site where REST inhibits PGR binding on *Esr1* locus. Arrows indicate PGR recruitment in the absence of REST in conditional knockout mouse uterus.

**Figure 4:**
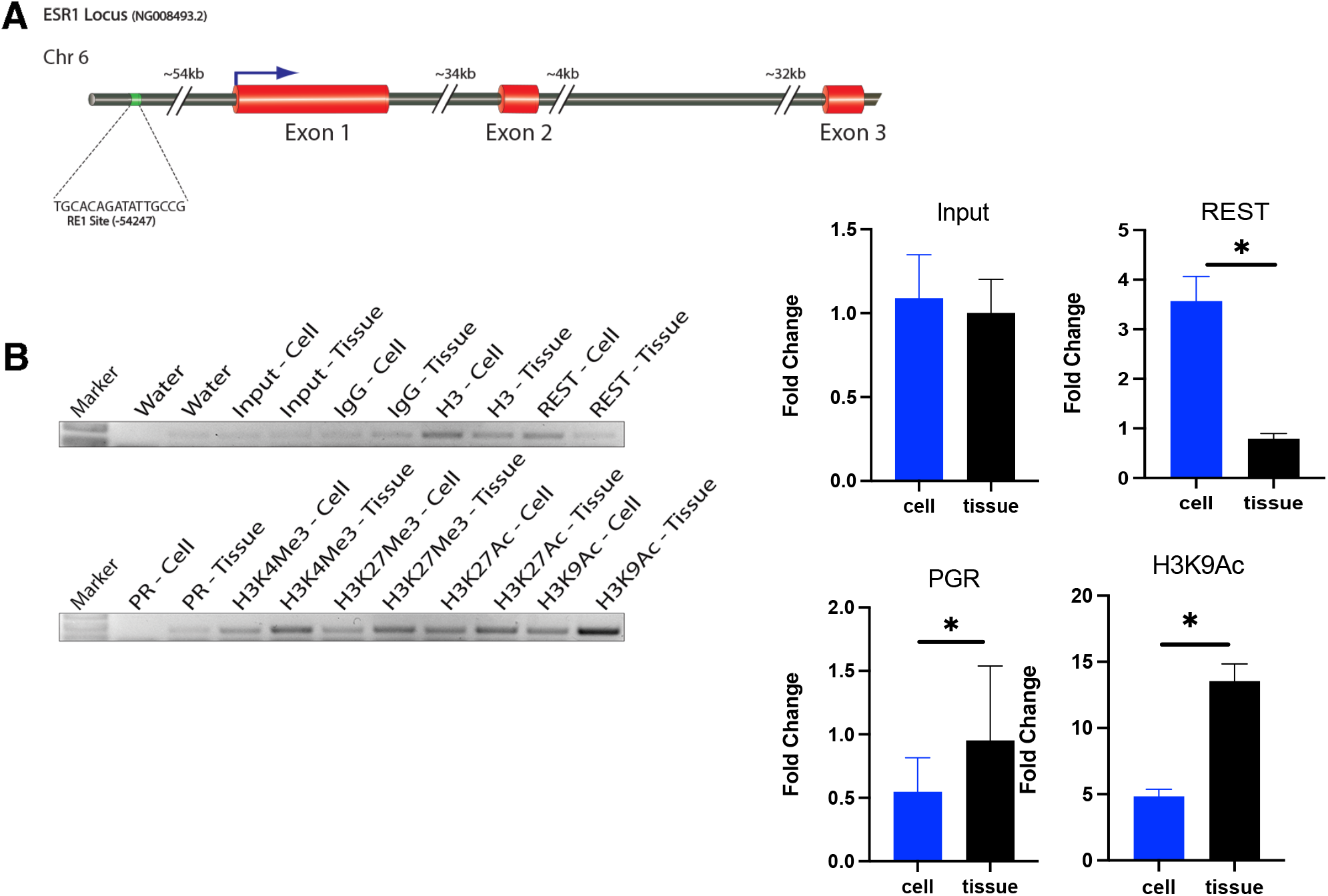
REST Regulates PGR Binding of *ESR1* Locus and Alters Histone Modifications. **(A)** Schematic of *RE-1* site of interest on *ESR1* locus. **(B)** ChIP-PCR binding levels in leiomyoma tissue versus cultured cells.

**Figure 5:**
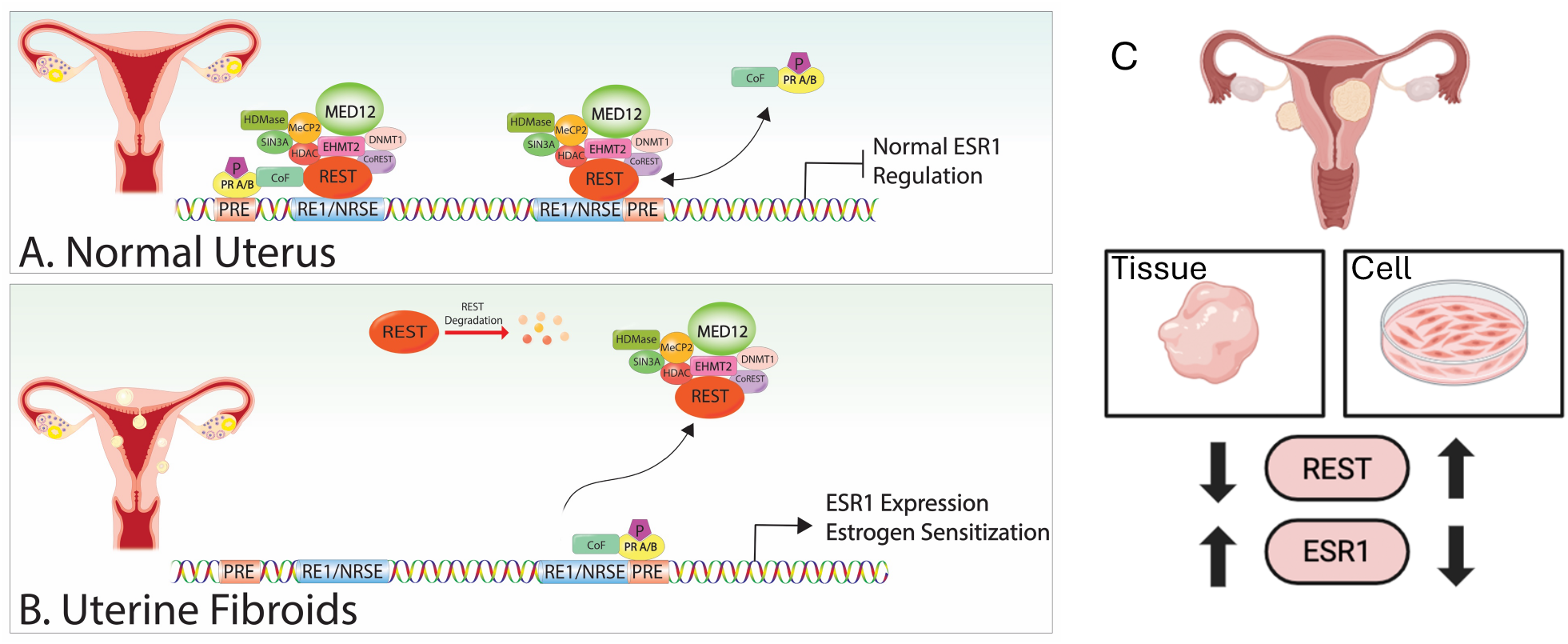
Schematic of proposed role of REST and PGR on *ESR1* regulation under (A) normal uterus, (B), uterine fibroids, and (C), schematic showing repression of ESR1 by increased REST in cell culture. Panel C was created using BioRender.com

We identified specific sites within the mouse *Esr1* locus where there was a difference in level of REST binding to *Esr1* between the control and cKO mice (Supplemental Figure 2A). REST binding sites of interest were prioritized using the presence of a *RE-1* motif and overlapping of *Esr1* locus as criteria. One of the sites of interest demonstrated adjacent and partial overlap in REST and PGR binding within 200 base pairs of each other on the *Esr1* locus. In REST cKO uterus, this site showed abolishment of PGR binding (Figure 3C). Conversely, an additional site showed less REST binding in the cKO but more PGR binding overlapping with the REST binding peak on the *Esr1* locus showing an inverse correlation between REST and PGR binding (Figure 3D).

### REST Regulates PGR Binding of *ESR1* Locus and Alters Histone Modifications

Using the UCSC Genome Browser (RRID:SCR_005780), the mouse genome was aligned with the human genome to determine sites of interest conserved in the human (Supplemental Figure 2B). In order to understand the epigenetic silencing of estrogen receptor in cultured leiomyoma cells, we investigated the epigenetic regulation of conserved *RE-1* sites within leiomyoma tissue and primary cells. We identified the conserved upstream *RE-1* site, which had shown an inverse correlation between REST and PGR binding in mouse, within the known *ESR1* regulatory region. This region, in addition to showing consistent REST and PGR binding as observed in mice, exhibited corresponding epigenetic changes within patient samples (Figure 4A) [23]. ChIP-PCR analysis revealed significantly low REST binding in leiomyoma tissue, reflecting low REST expression in UL, compared to cultured leiomyoma cells, representing higher REST expression during cell culture (Figure 4B). However, PGR binding and acetylated H3K9 levels were significantly higher in leiomyoma tissue compared to leiomyoma cells cultured from the same tissue (Figure 4B). Histone modification H3K4Me3 was also higher in leiomyoma tissue, however, it did not reach a level of significance compared to cultured cells. Collectively, these results suggest epigenetic changes that promote activation of transcription of *ESR1* in the absence of REST in leiomyoma tissue. Accompanying the robust expression of REST during cell culture, leiomyoma cells seem to acquire a repressive epigenetic signature, resulting in the silencing of *ESR1* expression.

## Discussion

It is well understood there is increased estrogen receptor expression and sensitivity which promote leiomyoma development [13, 15]. However, the mechanisms by which there is increased estrogen receptor expression and sensitization remain unclear. Here we provide a mechanism involving the loss of REST that promotes increased ESR1 expression in leiomyomas. We provide evidence that REST plays a role in epigenetic alterations of *ESR1* regulatory regions. Additionally, we demonstrate upstream REST regulator, PRICKLE1, acts through REST to alter ESR1 expression. We propose a mechanism where cultured myometrial and leiomyoma cells lose ESR1 expression due to increased REST expression and loss of its normal epigenetic control. We provide evidence that histone modifications related to *RE-1* binding sites in the *ESR1* locus are different between matched patient tissue and cultured cells, altering *ESR1* transcription levels.

Ablation of *Rest* in the whole uterus (*Rest*^*f/f*^*PR*^*+/Cre*^) as well as ablation of *Rest* in uterine mesenchymal cells (*Rest*^*f/f*^*Amhr2*^*+/Cre*^) resulted in a robust increase in *Esr1* expression (Figure 1C). This reflected the increased expression of *ESR1* seen in human uterine leiomyoma (Figure 1A). While it has been reported there is increased ESR1 expression in leiomyoma, there is no prior animal model that reflects estrogen receptor expression or increased estrogen signaling seen in uterine leiomyoma. Animal models targeting *Med12* and *Wnt/β-catenin* generate a leiomyoma phenotype, but there is no direct evidence these represent increased estrogen receptor expression or sensitivity [24, 25].

We wanted to see if we could recapitulate the regulation of expression of estrogen receptor by REST *in vitro*. Our results showing increased ESR1 expression under gene knockdown as well as CRISPR-Cas9 editing demonstrate that we can capture REST’s regulation of ESR1 seen *in vivo* also *in vitro* (Figure 2A, 2D). We also demonstrate PRICKLE1 acting upstream of REST, resulting in decreased REST expression thus leading to increased ESR1 expression (Figure 2B). There has been limited progress in the field of UL research to culture leiomyoma cells in a system which mimics *in vivo* patterns. We believe this may be, at least in part, due to REST returning in culture (Figure 2C). We believe modulation of REST could serve as a potential avenue to culture cells *in vitro* to recapitulate ESR1 expression, otherwise normally lost.

Our uterine gene epigenetic regulation data highlights important binding alterations around the promoter of *ESR1*. Many of the peaks which pass the cutoff criteria do not contain the consensus sequence for REST under the REST antibody condition (Table 3). It may be that the REST consensus is not yet well-defined. REST also may be binding globally through the mediation of other transcription factors. REST is already known to recruit other transcription regulatory complexes including CoREST, mSin3, SWI/SNF, PRC1, and PRC2 [26-28]. There is a need for further investigation of this discrepancy between peaks present with and without the consensus sequence. More genes involved in uterine gene epigenetic regulation controlled by REST could be studied in the future.

We investigated direct binding of REST to the *Esr1* locus and identified *RE-1* sites from the mouse which were conserved in human. Sites of interest were investigated both according to the mouse genome and conservation in human genome. Direct roles for the regulation of uterine transcriptome, cistrome, and chromatin dynamics by PGR have been investigated extensively [29-31]. It is well-established PGR does not act singularly as an activator or repressor [32]. We observed such phenomena in our data. One of the conserved *RE-1* sites within the *Esr1* locus that showed robust REST and PGR binding in the control group, lost PGR binding in the absence of REST (Figure 3C). We believe this suggests REST is needed for PGR binding, and REST may be recruiting co-repressors to enable PGR to act as a repressor of *Esr1* expression. This contrasts with an additional site of interest. In the cKO mouse and leiomyoma tissue, we see similar trends of less REST binding and more PGR binding at this site (Figure 3D, 4B). We believe this site may be acting as an activator/enhancer site for PGR on *ESR1* expression in the absence of REST. Both sites, one where PGR cooperates with REST to repress ESR1 as well as the one where PGR competes with REST for binding to an enhancer site, together will aberrantly maintain high ESR1 in UL in the absence of REST. We had previously reported proximal binding of REST and PGR on a vast majority of PGR targets showing an interaction between REST and PGR in the myometrium [15]. This demonstrated a novel regulatory role of REST on PGR mediated transcription Further studies are needed to analyze the functional importance of these *RE-1* sites separately.

It is recognized myometrium and leiomyoma smooth muscle cells silence ESR1 expression in culture. We hypothesized if the loss of REST is playing a role in the increased expression of ESR1 in UL, culturing leiomyoma cells will result in increased REST expression and specific repression of ESR1. Indeed, we observed a robust increase in REST protein expression in leiomyoma cells as well as increased binding of REST to *ESR1* locus (Figure 2A, Figure 4B). Additionally, we observed a significant decrease in H3K9Ac in the chromatin encompassing this *RE-1* site upon cell culture (Figure 4B). We also found trends in decreased H3K27Ac and H3K4Me3 in cultured cells, collectively showing increased recruitment of REST and silencing of *ESR1* expression upon cell culture.

Research of uterine fibroids suggests leiomyoma specific histone modifications such as H3K27Me3 and H3K4Me3 [7]. Additionally, Carbajo-Garcia et al., found a distinct clustering of H3K27 acetylation patterns in leiomyoma versus normal myometrium [33]. Leiomyoma were characterized by hyperacetylation of oncogenes and hypoacetylation of tumor suppressors. Reversal of the H3K27Ac modification resulted in the upregulation of tumor suppressor genes, otherwise downregulated in leiomyoma [33]. Yang et al. found Class I HDACs to be upregulated in leiomyosarcoma compared to adjacent myometrial tissue [34]. A similar pattern was observed in leiomyosarcoma cell lines, and inhibition of the HDACs decreased cell proliferation. Interestingly, Class I HDAC protein levels increased from benign fibroid tissue to malignant leiomyosarcoma [34]. REST has also been found to recruit histone H3K9 methyltransferase, G9a as well as MED12, which is frequently mutated in UL [35, 36]. Collectively, these results suggest a mechanistic role for histone modifications resulting from the loss of REST in the development of uterine fibroids. We provide novel evidence regarding the role of REST on histone modification changes in the *ESR1* locus involved in leiomyoma development. Future research is warranted to determine how targeted mutations in *RE-1* binding sites within the *ESR1* locus may affect leiomyoma histone modifications and *ESR1* expression.

There has been a large effort to develop leiomyoma cell culture models which recapitulate hormone response seen in leiomyoma pathophysiology [37, 38]. There has been the establishment of 3D spheroid systems as well as immortalization of leiomyoma and myometrial cells, however, there is a lack of optimization for all cell culture models. More specifically, leiomyoma cell culture methods fail to recapitulate ESR1 expression levels to the like of leiomyoma present in the body [37, 39]. Our results could lead to the development of a novel cell culture system which restores ESR1 expression through PRICKLE1 and REST regulation. Collectively, our results which mirror the dysregulation of ESR1 expression in UL, strongly support the relevance of *Rest* cKO animal model to the study of steroid hormone pathways in uterine fibroids.

Current uterine fibroid medications utilize progesterone and estrogen receptors as targets. Medications specifically targeting progesterone, known as selective progesterone receptor modulators (SPRMS), pose concerns surrounding long-term usage, at least in part due to gene activator and repressor functions of PGR within the same tissue [10-12]. Our results also suggest a context dependent function of PGR in the regulation of *ESR1* expression. Despite attempts to determine effective medications targeting estrogen, challenges remain. Aromatase inhibitors can result in fibroid shrinkage but leave patients with a myriad of side effects mimicking menopause including hot flashes and osteoporosis. Selective estrogen receptor modulators (SERMs) such as tamoxifen and raloxifene have been investigated as well. Tamoxifen was ineffective at reducing fibroid size and lead to ovarian cyst development in some patients. Similarly, raloxifene was ineffective at reducing fibroid size substantially and demonstrated poor pharmacokinetic specificity [13]. We believe this study reveals a novel understanding of increased estrogen sensitivity in leiomyoma pathophysiology and will aid in the development of better endocrine therapies for uterine fibroids.

## Methods

### Study Approval

Studies involving patient derived hysterectomy specimen were approved by and performed in accordance with the University of Kansas Medical Center Policies and Procedures Relating to Human Subjects (IRB# 00160213). All patients gave informed consent when donating their uterine tissue samples to the University of Kansas Medical Center. All mouse experiments were approved by the University of Kansas Medical Center IACUC protocols (23-02-296) and adhere to NIH guidelines for care and use of laboratory animals.

### Tissue Collection and Cell Culture

Matched myometrium and leiomyoma tissue samples were obtained from pre-menopausal women undergoing hysterectomies at the University of Kansas Hospital (Kansas City, KS). Criteria for patients in this study exclude women undergoing a hysterectomy for a primary condition other than uterine leiomyoma and those taking hormone therapy in the three months preceding surgery. Myometrial primary cells were prepared from samples as previously described [16].

### ChIP-Seq Analysis

An unbiased whole genome profiling of REST and PGR binding was obtained by chromatin immunoprecipitation followed by massively parallel sequencing (ChIP-Seq). ChIP-Seq data was generated from REST-ChIP in control (*Rest*^*fl/fl*^), REST-ChIP in *Rest* conditional knockout (*Rest*^*fl/fl*^ *PR*^*+/Cre*^), PR-ChIP in control (*Rest*^*fl/fl*^), and PR-ChIP in *Rest* conditional knockout (*Rest*^*fl/fl*^ *PR*^*+/Cre*^) samples. We used corresponding input DNA controls from WT and KO samples to measure total chromatin before IP. ChIP-Sequencing was performed in an Illumina NovaSeq 6000 sequencing machine (Illumina, San Diego, CA) at a 100-base paired-end read resolution. After read quality assessment (FastQC [40]) and adapter trimming (Cutadapt [41]), the sequenced reads were mapped to the mouse reference genome (GRCm38) using the Bowtie2 [42] software. Sequencing produced between 31.5 and 56.5 million reads per sample of which 97% to 99% mapped to the reference genome. Significantly enriched ChIP regions (relative to the corresponding Input sample) were detected using the Model-based Analysis of ChIP-Seq (MACS2 [43]) software. The analysis was run under default settings in the paired-end mode with the q-value cutoff for calling significant regions set at 0.05. The peak regions were scanned for *RE-1* and *Pgre* binding motifs with a position weight matrix (PWM) similarity cutoff set at 70%. The number of significantly enriched regions (enrichment FC ≥ 1.5, q-value ≤ 0.05) detected for each antibody is given in Figure 4C. The mapped regions were visualized using the Integrative Genome Viewer (IGV [44]). A motif analysis for known transcription factor binding sites in the enriched regions was performed using the HOMER [45] software. The heatmaps of sequencing-depth-normalized read coverage around 3 kb up-stream and down-stream from the transcription start site (TSS) of all genes were generated using the deepTools [46] software.

### ChIP-PCR

Using ChIP-seq analysis, we identified several sites within the mouse *Esr1* locus where there was a significant difference in level of REST binding to *Esr1* between the control and cKO mice. Sites of interest were prioritized using the presence of a *RE-1* motif and overlapping of *Esr1* gene as criteria. Using the UCSC Genome Browser (RRID:SCR_005780), the mouse genome was then aligned with the human genome to determine *RE-1* sites conserved in the human. Primer sets were designed (Table 2) for human sites of interest according to standard primer design protocol. ChIP was performed for analogous patient cell and tissue samples. Patient primary leiomyoma cells were grown in antibiotic free media (Thermofisher Scientific, DMEM #11960-044) supplemented with fetal bovine serum, bovine calf serum, L-glutamine, and penicillin-streptomycin in a 37°C 5% CO_2_ incubator. Corresponding patient fibroid tissue was biopulverized using RPI Corp 141275 Biopulverizer (Mount Prospect, IL). Once the cells reached 100% confluency, both the cells and biopulverized tissue underwent standard ChIP kit protocol (Cell Signaling, #14209S, Cell Signaling Technology RRID:SCR_002071). Antibodies used for ChIP included rabbit IgG and H3XP as controls (Cell Signaling Technology Cat# 2729, RRID:AB 1031062, Cat# 4620, RRID:AB_1904005). Antibodies included Anti-REST, Anti-PR, Anti-H3K4Me3, Anti-H3K27Me3, Anti-H3K27Ac, and Anti-H3K9Ac (Millipore Cat# 07-579, RRID:AB_310728, Thermo Fisher Scientific Cat# MA1-12626, RRID:AB_1086828, Cell Signaling Technology Cat# 9751, RRID:AB_2616028, Cat# 9733, RRID:AB_2616029, Cat# 8173, RRID:AB_10949503, Cat# 9649, RRID:AB_823528). The samples underwent standard PCR and run on a 1.5% agarose ethidium bromide gel before being imaged using a GelDoc imager.

**Table 1.**
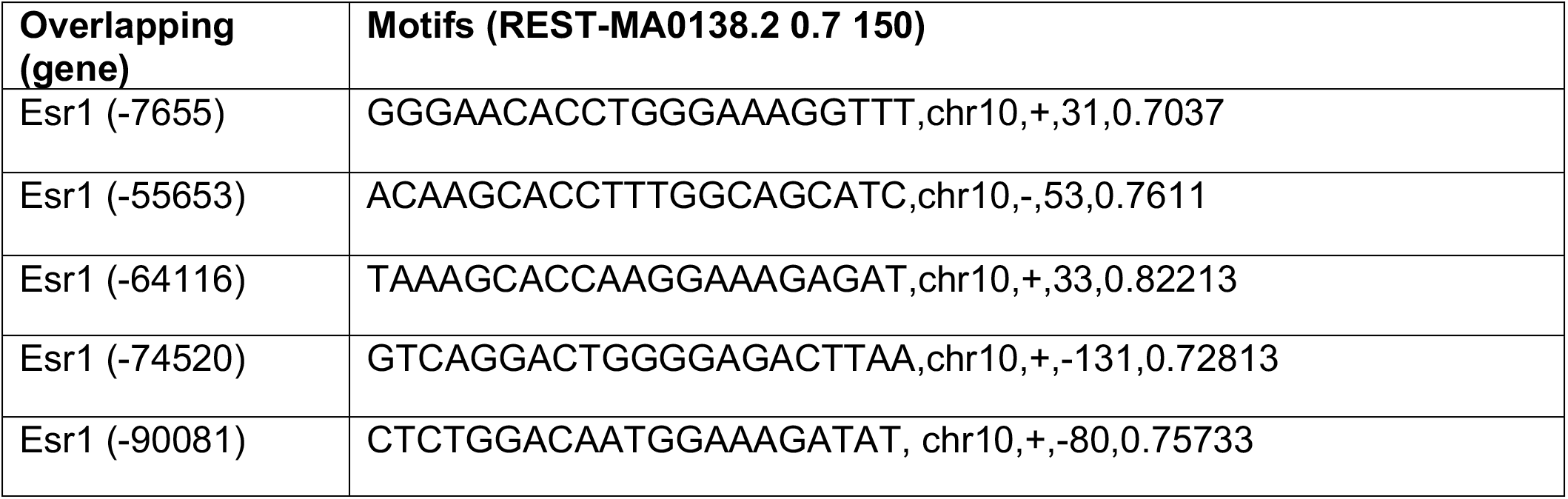
*RE-1* motif binding sites within *Esr1* locus in ChIP-seq data of *Rest*^*fl/fl*^ *PR*^*+/Cre*^ mouse model.

**Table 2.**
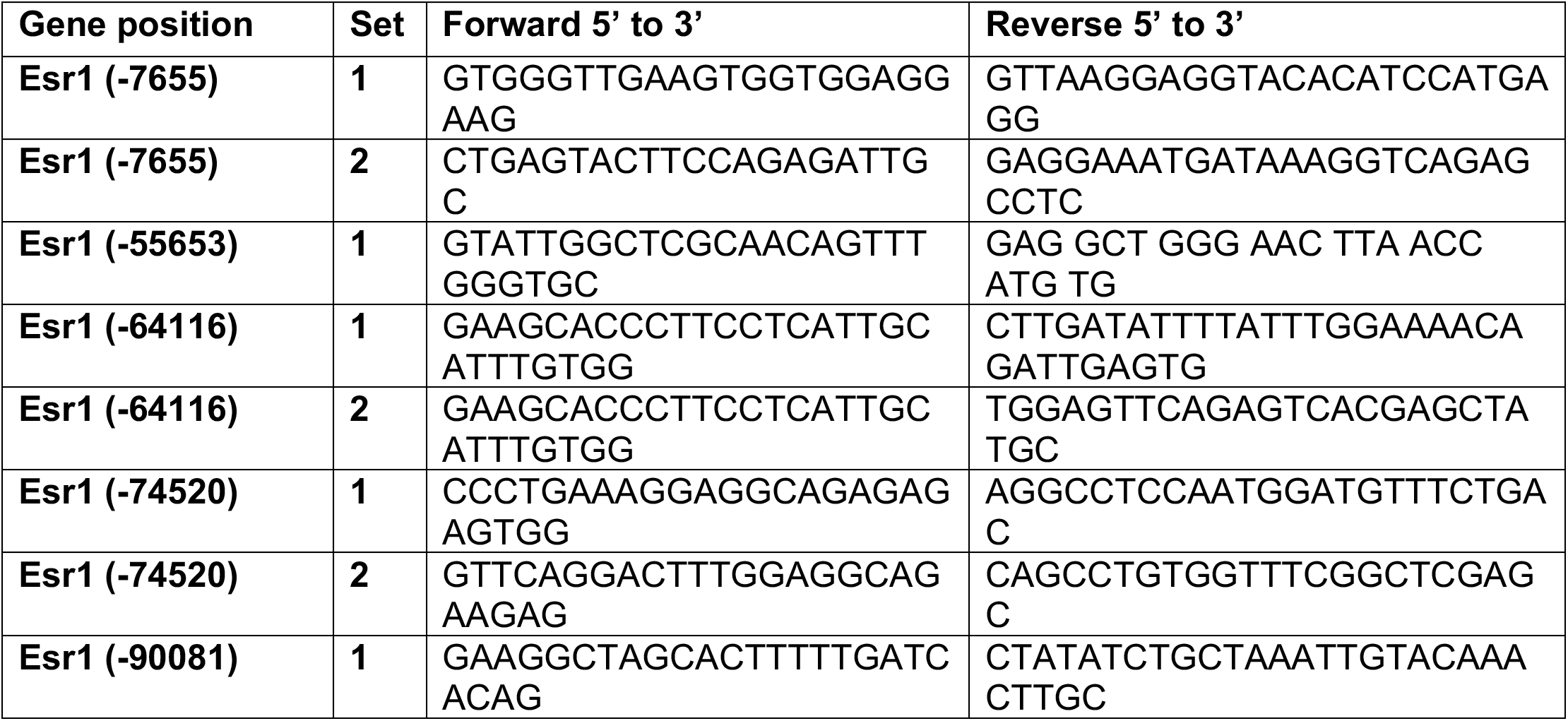
Primer design for REST binding sites within *ESR1* locus conserved in human genome.

**Table 3:** Antibody efficiency of REST and PGR in *Rest*^*fl/fl*^ and *Rest*^*fl/fl*^ *PR*^*+/Cre*^.

| REST Antibody condition | # Peaks pass cutoff | # Peaks with *RE-1* consensus site |
| --- | --- | --- |
| *Rest<sup>fl/fl</sup>* | 121,768 | 26,929 |
| *Rest<sup>fl/fl</sup> PR<sup>+/Cre</sup>* | 58,587 | 12,645 |
| PGR Antibody condition | # Peaks pass cutoff | # Peaks with *Pgre* consensus site |
| *Rest<sup>fl/fl</sup>* | 5,442 | 5,349 |
| *Rest<sup>fl/fl</sup> PR<sup>+/Cre</sup>* | 1,724 | 1,688 |

### Transfections

Cultured myometrial cells were plated in 6-well plates at 200,000 cells per well in antibiotic free media (Thermofisher Scientific, DMEM #11960-044). After 24 hours of incubation at 37°C 5% CO_2_, cells were transfected with 100pmol of Silencer Select siRNA *REST* complexes (Ambion, #4392421) using Lipofectamine 2000 transfection reagent (Invitrogen, Cat#11668019). Similarly, transfections investigating *PRICKLE1* used *PRICKLE1* Silencer Select siRNAs (IDT, #170291100,170291101,170291102). Control wells included negative control DsiRNA (DS NC-1, IDT, #51-01-14-04). The plate incubated at 37°C 5% CO_2_ with the complexes for 48 hours. At the conclusion of the 48 hours, protein was collected from each well.

### Protein Extraction and Western Blotting

Cells were lysed in 1x cell lysis buffer (Promega, Cat#E153A) and supplemented with protease and phosphatase inhibitor cocktails (Sigma-Aldrich Co., Cat# P0044, P8340). Each tube of protein collected underwent 20 seconds of sonication (Fisherbrand Model 120 Sonic Dismembrator) at 40% amplitude followed by 10 seconds on ice for three rounds. Each tube was then centrifuged at 20,000g for 15 minutes at 4°C. Protein was quantified using Bio-Rad Protein Assay (Bio-Rad, Cat#5000006). Proteins were detected using SuperSignal West Femto Maximum Sensitivity Substrate (ThermoFisher Scientific, Cat# 34096) and blots were imaged using BioRad ChemiDoc MP (Bio-Rad Laboratories RRID:SCR_008426). Antibodies used for western blotting included Anti-PRICKLE1, Anti-REST, Anti-ESR1, Anti-Mouse IgG HRP Conjugate, and Anti-Rabbit IgG HRP Conjugate (Proteintech Cat# 22589-1-AP, RRID:AB_2879129, Millipore Cat# 07-579, RRID:AB_310728, Cell Signaling Technology Cat# 13258, RRID:AB_2632959, Promega Cat# W4021, RRID:AB_430834, Cat# W4011, RRID:AB_430833). Protein loaded was quantified using Anti-Beta-actin (GenScript Cat# A01546, RRID:AB_1968817). Western blot protein bands were quantified using ImageJ software. All antibodies were validated by gene knockdown or knockout.

### Histology and Staining

Uterine tissues were fixed in 4% paraformaldehyde for 24 hours and then processed for paraffin embedding. Tissues were deparaffinized in xylene and rehydrated in a series of ethanol washes and stained for immunofluorescence. For immunofluorescence staining, antigen retrieval was performed following rehydration by boiling in antigen unmasking solution (Vector Labs, Cat# H-3301). Following antigen retrieval, tissues were stained with primary antibody anti-ESR1 (Cell Signaling Technology Cat# 13258, RRID:AB_2632959, 1:100). Nuclei were stained with DAPI (Invitrogen, Cat# D1306). Alexa Fluor 488 secondary antibody was used for all immunofluorescence staining (Jackson ImmunoResearch Labs Cat# 711-545-152, RRID:AB_2313584, 1:200). All histological samples were imaged using a Nikon Eclipse 90i fluorescent microscope with Nikon Digital Sight color DS-Fi1 camera.

### Taqman qRT-PCR

Total RNA was isolated from uterine tissue samples stored in RNAlater (Invitrogen, Cat# AM7020), biopulverized, and placed into TRIzol Reagent (Invitrogen, Cat# 15596026). Following TRIzol, a series of chloroform, isopropanol, and ethanol washes were used to isolate the RNA. Following quantification using Nanodrop spectrophotometer, aliquots of RNA were reverse transcribed using a High Capacity cDNA Reverse Transcription Kit (Applied Biosystems, 4368814, Applied Biosystems RRID:SCR_005039). TaqMan assay for *ESR1* (IDT, Hs.PT.58.14846478) was used to quantify gene expression difference utilizing the ΔΔC(T) method with housekeeping gene *RNA18S* (IDT, Hs.PT.39a.22214856.g) for myometrial and leiomyoma human tissue.

### Luciferase Assay

Myometrial cells were cultured and transfected as described above. The control group consisted of cultured MCF7 cells plated as previously described [47]. Each well received ERE-TAL-Puro TF Reporter Lentivirus (LipExoGen, LTV-0046-3S) at an MOI of 1. The 6-well plate incubated at 37°C 5% CO_2_ for 48 hours. At the conclusion of 48 hours, the cells were trypsinized and replated in a 96-well opaque black plate according to the Firefly Luc One-Step Glow Assay Kit protocol (ThermoScientific, #16196, #16197). Each group was plated for quadruplicates. The 96-well plate incubated at 37°C 5% CO_2_ for 24 hours to allow for cell attachment. After 24 hours, each experimental group had one set of quadruplicates receive charcoal stripped media and another receive 10 nM of E_2_ in charcoal stripped media. The plate incubated at 37°C 5% CO_2_ for 48 hours. After 48 hours, a Firefly Luc One-Step Glow Assay was performed according to the Pierce Firefly Luc One-Step Glow Assay Kit protocol (ThermoScientific, #16196, #16197). The plate underwent a top read on a luminometer with an integration time of 500 ms.

### Statistical analysis

Leiomyoma samples were compared to myometrial controls from the same patient. For animal studies, *Rest* cKO mice uterine samples were compared to age-matched control mice at the same stage of the estrous cycle. Quantitative experiments were repeated with at least three independent biological replicates. Sample size was determined using desired power of 80%, fold change of 2, and p<0.05. Statistical significance was determined using students T-test to determine change from the control sample.

## Acknowledgements

The authors acknowledge the KIDDRC (NIH U54 HD 090216) for guidance with imaging at the University of Kansas Medical Center, Kansas City, KS, 66160.

