## Supplemental Data for "Regulatory Role for Tumor Suppressor REST on Estrogen Receptor (*ESR1*) Expression and Leiomyoma Pathophysiology"

**CRISPR/Cas9 Editing**

HEK293T cells were maintained in media (DMEM #11995-065) supplemented with 10% FBS, 1% pen/strep, and 1% L-glutamine at 37°C 5% $CO_2$. The cells in 6-well plates were incubated with a transfection mixture for 24 hours at 37°C. The transfection mixture consisted of 100μL JetPrime transfection buffer/mL of media (PolyPlus #712-60), 10μL of Ready-to-Use Lentiviral Packaging Plasmid Mix (Cellecta CPCP-K2A), and 2μL CRISPR/Cas9 plasmid (GenScript REST CRISPR Guide RNA 1 in pLentiCRISPR v2 U3449CJ090-1/S78294, GenScript REST CRISPR Guide RNA 2 in pLentiCRISPR v2, U3449CJ090-2/S78296). The transfection mixture was briefly vortexed, spun down, and kept on ice. Two μL JetPrime transfection reagent/mL (Polyplus #114-07) of media was then added to the transfection mixture followed by a brief vortex and spin down. The mixture was then left at room temperature for 10 minutes. After 10 minutes, the transfection mixture was added to the plate of cells and left to incubate for 30 minutes at 37°C. The plate was then removed from the incubator, gently swirled for 30 seconds, and returned to the incubator. This was repeated for 2 hours. At 24 hours post transfection, media was changed, and cells were left to incubate for another 24 hours at 37°C.To harvest the virus, the media was removed with a plastic syringe and filtered through a sterile 0.22μM pore size PES membrane filter into a conical tube. Twice the amount of media volume was added back onto the plate. The plate was again left to incubate at 37°C. Harvesting of the virus was repeated at 72 hours post transfection. Once filtered, the virus was pipetted in 1 ml aliquots into sterile Eppendorf tubes and spun down in 4°C at max speed for 1-2 hours in 30-minute increments. At the conclusion on the spin down, a pipette was used to remove and dispose of 900μL from the top of each tube. The remaining 100μL left in each tube was shaken on a vortex with an Eppendorf rack/attachment for at least 30 minutes. To perform the infection, 500μL of media was added to one Eppendorf tube. The viral suspension was taken up and added to each subsequent tube, pipetting up and down gently until all the virus was in a single tube. Immortalized leiomyoma cells were infected at a confluency of 50-60% on a 3 cm plate. A volume of 1.5 mL of viral suspension was used, and cells were left to incubate at 37°C. Infection was repeated 2-3 times. After the final infection, cells were split 24 hours post-infection if they reached confluency or were overconfluent. Selection started at least 48 hours post-infection. Puromycin was gradually increased to 1:1000 dilution of 5μg/μL. CRISPR cells underwent 7-8 passages or 21 days of selection. Immortalized uninfected leiomyoma cells were used as a control group during selection. Some cells were used for western blotting, and the rest were used for a single colony assay. A single colony assay was performed using a 96-well plate. Chosen colonies originated from a single-cell-based colony. Media was changed every 2-3 days. The plate was incubated at 37°C, and clones were detectable by microscopy after 4 to 5 days. We selected 4 single colonies and screened them by western blots at passage 3 and 6. We ran Taqman qRT-PCR at passage 3 and passage 6 to confirm knockout of *REST* (IDT, Hs.PT.58.24545780). In addition, housekeeping gene *RNA18S* (IDT, Hs.PT.39a.22214856.g) and *REST* targets, *EZH2* (Hs.PT.58.38882546), *ESR1* (Hs.PT.58.14846478), *PGR* (Hs.PT.58.3493960), *COL3A1*(Hs.PT.58.4249241), and *COL1A2* (Hs.PT.58.26714610) were investigated by Taqman qRT-PCR.

**Supplemental Figures**

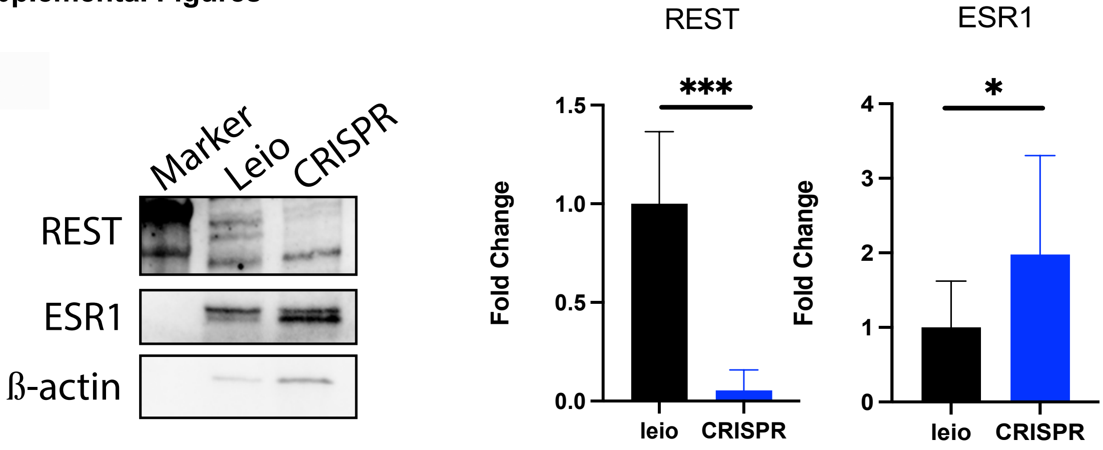


**Supplemental Figure 1:** Western blot of immortalized leiomyoma (DDHLM) cell line with complete knockout of *REST* using CRISPR-Cas9 lentivirus.

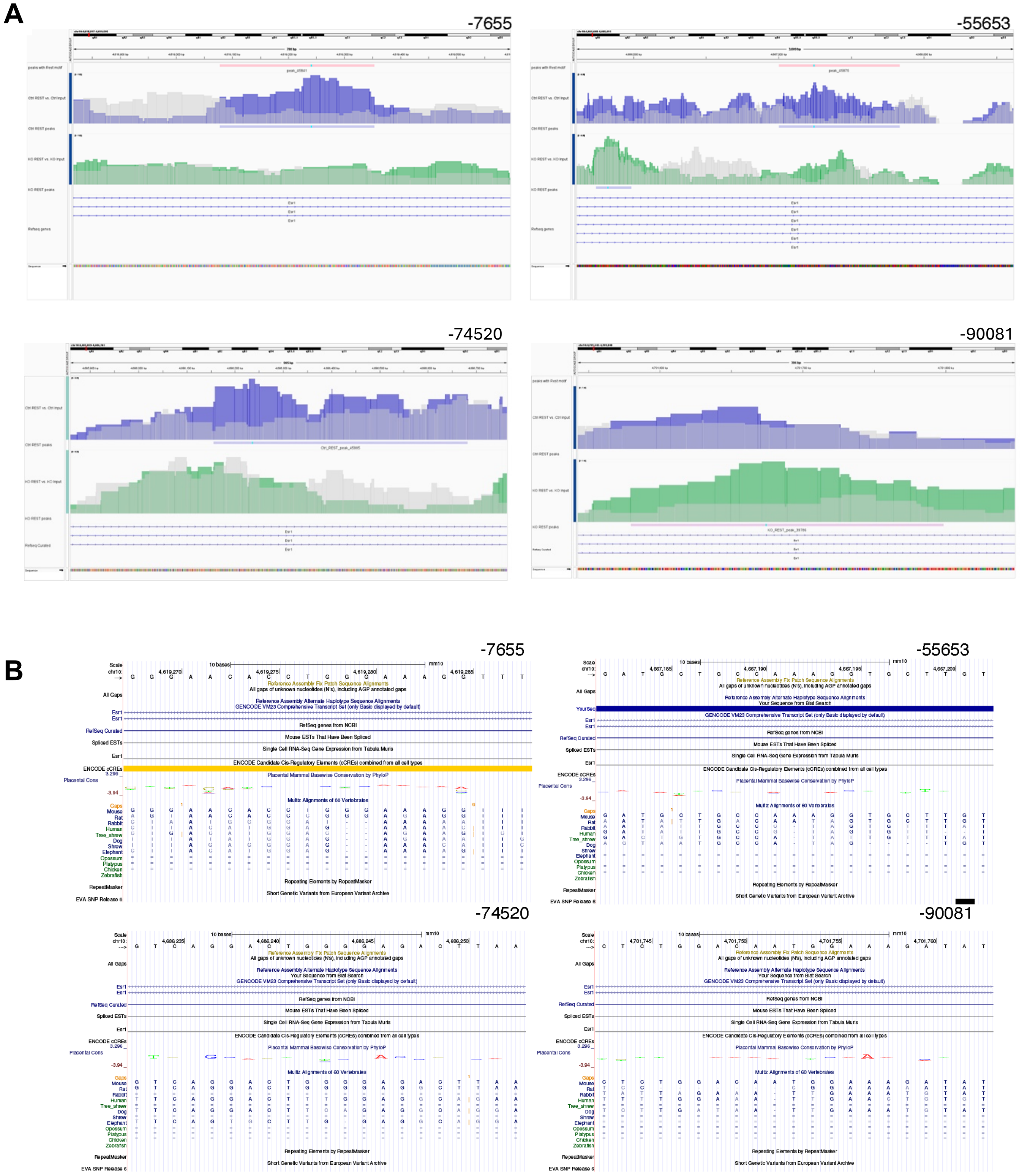


**Supplemental Figure 2: *RE-1* Site Analysis within *ESR1* Locus. (A)** Binding peaks of *RE-1* sites on *Esr1* locus (blue) against input background binding (grey). **(B)** Alignments of *RE-1* sites on *Esr1* locus in mouse genome against the *ESR1* locus in human genome.
